# Comparing multiscale, presence-only habitat suitability models created with structured survey data and community science data for a rare warbler species at the southern range margin

**DOI:** 10.1101/2022.09.21.508867

**Authors:** Lauren E. Whitenack, Sara Snell Taylor, Aimee Tomcho, Allen H. Hurlbert

## Abstract

Golden-winged Warblers (*Vermivora chrysoptera*, Parulidae) are declining migrant songbirds that breed in the Great Lakes and Appalachian regions of North America. Within their breeding range, Golden-winged Warblers are found in early successional habitats adjacent to mature hardwood forest, and previous work has found that Golden-winged Warbler habitat preferences are scale-dependent. Golden-winged Warbler Working Group management recommendations were written to apply to large regions of the breeding range, but there may be localized differences in both habitat availability and preferences. Rapid declines at the southernmost extent of their breeding range in Western North Carolina and Georgia necessitate investigation into landscape characteristics governing distribution in this subregion. We describe patterns of Golden-winged Warbler presence in Western North Carolina by examining habitat variables at multiple spatial scales using data from standardized Audubon North Carolina (NC) surveys and unstructured community scientist checklists submitted to eBird. We compared model performance and predictions between Audubon NC and eBird models and found that Golden-winged Warbler presence is associated with sites which, at a local scale (150m), have more young forest, less mature forest, and more road cover, and at a landscape scale (2.5km), have less road cover. eBird and Audubon models had similar parameter estimates for all of the land cover variables and similar overall performance. Based on parameter estimates, road density at the landscape scale (2.5km) is the primary variable driving the difference between Golden-winged Warbler breeding sites and random background sites in Western North Carolina. Our results show that eBird data can produce species distribution modeling results that are similar to results obtained from more standardized survey methods.

## Introduction

Habitat loss is a primary threat to biodiversity in the present day [1,2]. Migratory birds may be especially vulnerable to habitat loss since they rely on the persistence of multiple quality habitats for breeding, stopover, and wintering [3,4]. As such, understanding the habitat associations of migratory birds can help predict patterns of distribution and abundance, and inform management practices and conservation efforts. The Golden-winged Warbler (*Vermivora chrysoptera*, Parulidae) is a migrant songbird which has been declining at an average of 1.85% per year for over 50 years across their range [5]. Due to its vulnerable status, the Golden-winged Warbler has been assigned to the Partners in Flight Red Watch List [6], the United States Fish and Wildlife Service’s list of Birds of Conservation Concern [7] and is a candidate for listing under the Endangered Species Act [8].

Currently, Golden-winged Warblers breed in two main regions: the Great Lakes region of southeastern Canada and the northern-midwestern United States, and the Appalachian region in select moderate-to-high elevation sites in the Appalachian Mountains [9]. Within their breeding range, Golden-winged Warblers are found in early successional habitat near mature hardwood forest [10,11]. During the breeding season, Golden-winged Warblers use multiple vegetation layers. Golden-winged Warblers build their nests on the ground in the herbaceous layer or just above the ground in shrubs [12,13]. Shrub cover provides protection from predators and surrounding mature hardwood forest is used for male perches, nesting material, and foraging ground [11,12,14,15]. Because of their complex habitat requirements, researchers have focused on studying Golden-winged Warbler breeding habitat associations at multiple spatial scales. Studies have shown that variables important at the scale of the nest site, such as percent herbaceous and shrub cover, differ from variables important at larger scales, such as percent mature forest [16,17,18]. Despite the general trends, Golden-winged Warbler breeding habitat associations vary between geographical areas and with landscape context [16,18]. Thus, it is important to study Golden-winged Warbler breeding habitat associations throughout their breeding range and at multiple spatial scales to understand these regional differences.

Golden-winged Warblers are especially vulnerable in the southern Appalachian Mountains at the southernmost extent of their breeding range. Data from the North American Breeding Bird Survey indicate that in Western North Carolina, Golden-winged Warbler populations have decreased by approximately 6.5% per year from 1993-2019 [5]. Habitat loss due to human development and maturation of early successional habitat, as well as brood parasitism by Brown-headed Cowbirds (*Molothrus ater*) have contributed to this decline [12]. In the Appalachian Region, Golden-winged Warblers hybridize extensively with Blue-winged Warblers (*Vermivora cyanoptera*), especially at moderate elevations [9]. Traditionally, it was thought that human-related habitat changes caused an increase in hybridization between the two species over the last 50 years. However, recent genomic work by Toews et al. (2016) suggests that the two *Vermivora* species have been hybridizing for millennia [19]. Toews et al. (2016) recommend that Golden-winged Warbler conservation efforts focus on maintaining the phenotypic diversity within the *Vermivora* complex, regardless of whether the two species are indeed distinct. Thus, declines in Golden-winged Warbler populations necessitate further research and understanding of the breeding ecology and habitat associations of this species, especially at the southern limit of its breeding range.

The Golden-winged Warbler Working Group (GWWG) was founded in 2003 to facilitate collaboration among scientists to produce best management practices for the conservation of Golden-winged Warblers throughout their breeding and wintering ranges [20]. The GWWG best management practices for the Appalachian Region are designed to apply to Golden-winged Warbler populations in the Appalachian Mountains from New York to Georgia. Because local Golden-winged Warbler habitat associations may vary within the large latitudinal gradient of the Appalachian Region, it is critical to examine how these management guidelines align with Golden-winged Warbler habitat associations across the region. Difficulty in identifying early successional habitat with appropriate granularity across large spatial scales has prevented large-scale quantitative analyses of habitat associations to support these recommendations. Fine-tuning these management recommendations based on localized differences in habitat associations could vastly improve conservation efforts of Golden-winged Warblers, especially at the limits of their breeding range.

For over thirty years, Audubon North Carolina (Audubon NC) has conducted playback surveys during breeding season to collect data on the abundance and distribution of breeding Golden-winged Warblers in Western North Carolina using the Golden-winged Warbler Atlas Project protocol [21]. Audubon NC survey locations are chosen based on drive-by habitat assessments, aerial photo review, proximity to known locations within dispersal range, and private landowner cooperation, which could focus survey effort on certain parts of the landscape while excluding others. Audubon NC surveys are conducted with the primary goal of finding new Golden-winged Warbler habitat, and much of what is known about the current distribution of Golden-winged Warblers in North Carolina can be attributed to Audubon NC and the Golden-winged Warbler Atlas Project. Analysis of the habitat associations of breeding Golden-winged Warblers in Western North Carolina could help identify parts of the landscape that may be suitable for breeding birds but have not been surveyed.

With the increase in popularity of the community science platform eBird, avian presence and abundance data is now freely available for scientists to use to study species’ ranges and track changes in distribution over time [22]. eBird users can submit bird observations at any location and time, resulting in over 70 million complete checklists worldwide, and over 1 million in North Carolina alone at the time of this publication [23]. Notably, eBird data are unstandardized and usually collected by non-professionals, potentially resulting in a noisier dataset due to misidentifications and differences in data collection methods. Unlike structured surveys such as the Golden-winged Warbler Atlas Project, eBird data are not collected with a specific conservation or scientific goal. Despite these shortcomings, eBird data are increasingly being used successfully to understand distributions and habitat associations of bird species [24–28]. Community science data require fewer resources to collect than more traditional survey methods, which require time and financial resources to organize and implement. Additionally, community science data such as eBird can be used to supplement or replace more structured survey data to increase the accuracy of distribution models and improve understanding of species’ habitat associations [25]. Since structured surveys require time and financial resources, it would be valuable to know if Golden-winged Warbler habitat models created with community science data produce similar results to those created with structured survey data.

Because both structured survey data (Audubon NC) and community science data (eBird) are abundant and available in the area, Western North Carolina is the ideal study site to compare habitat models created with these two datasets. Additionally, Western North Carolina is a subregion of considerable conservation concern, since it is located at the southernmost extent of the Golden-winged Warbler breeding range. Our goals in this paper are twofold: (1) to describe habitat associations of Golden-winged Warblers in Western North Carolina at multiple spatial scales, and (2) to determine whether a model created using eBird data would yield the same results as a model created with Audubon NC data.

## Methods

We studied Golden-winged Warbler habitat associations across Western North Carolina within and around previously established focal areas as defined by [20] and within the breeding range defined by the U.S. Geological Survey – Gap Analysis Project [29] (Fig 1).

**Fig 1.**
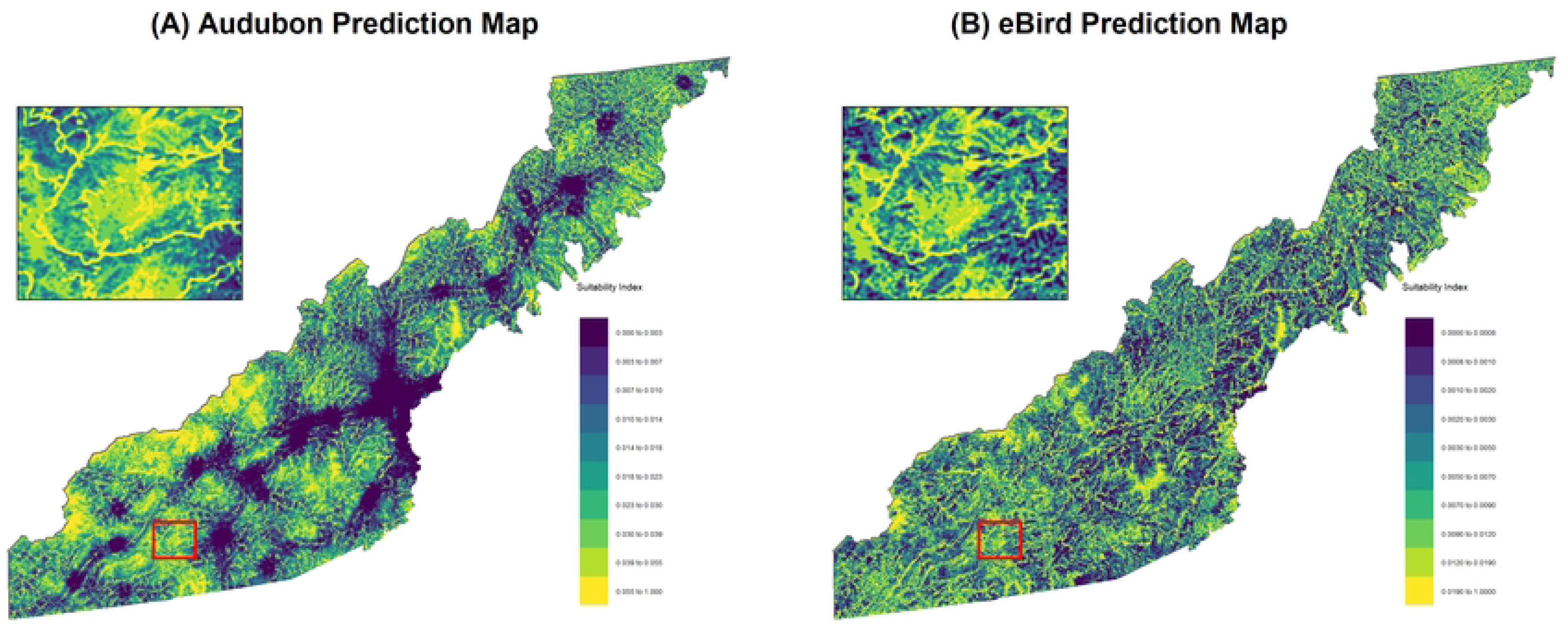
Study area in Western North Carolina, United States.

We conducted our analyses using two sets of Golden-winged Warbler presence data: Audubon NC survey data and community science data from eBird. First, we obtained Golden-winged Warbler presence data from Audubon NC collected during breeding season (May-July) from 2000-2020. The data from Audubon NC were collected using standardized playback surveys as outlined in the Golden-winged Warbler Field Survey Protocol prepared by the Cornell Lab of Ornithology [21]. Audubon NC surveys were conducted at sites that were determined by visual inspection to be potentially suitable, usually roadside or on private lands managed for Golden-winged Warblers with the permission of the landowner. Second, we downloaded Golden-winged Warbler observations from eBird during the breeding season from 2000-2020 [23]. We used stationary and incidental checklists only, excluding traveling checklists for which there is greater spatial uncertainty surrounding the precise location of target birds. We removed eBird checklists that were submitted under the Golden-winged Warbler Atlas Project protocol, since Audubon NC now submits their survey results to eBird.

To further correct for data redundancy, we searched both datasets for all presence locations within 100m of each location and kept only the location of the most recent Golden-winged Warbler observation. Notably, many known Golden-winged Warbler locations visited by eBirders may have been initially identified by Golden-winged Warbler Atlas Project surveys, and vice versa. To account for this overlap between datasets, we assigned each point to the dataset with the earliest record of a presence within 100m of that point, starting in 1988 when the Audubon NC surveys began. This redundancy analysis resulted in an Audubon sample size of 279 presence points and an eBird sample size of 86 presence points.

We obtained 2014 landcover data from the United States Forest Service LandFire Data Distribution Site with a pixel size of 30m by 30m [30]. We chose to use the 2014 Existing Vegetation Height (EVH) LandFire dataset since it includes forest height classification, unlike the National Land Cover Dataset (NLCD), which includes vegetation categories without vegetation height data.

Using R, we created two sets of rasters from each of the initial LandFire raster layers: one set describing percent land cover type within a 150m circular buffer of each pixel and the other describing percent land cover within a 2.5km circular buffer of each pixel [31]. Layers describing percent land cover type within a 150m buffer included percent forest of height 0-10 meters, percent forest of height 25-50m, percent herb/shrub, and percent road cover. Layers describing percent land cover type within a 2.5km buffer included percent forest of height 25-50m, percent road cover, and percent developed land. These landcover covariates were selected based on the management recommendations outlined by the GWWG [20].

We created separate models for Audubon NC and eBird datasets. For both datasets, we created 10,000 background points for model training by sampling from environment polygons that extended 10km around each presence point [32]. We extracted environmental variables at presence and background points in each dataset. We used generalized linear models with a logit link to determine how sites could be predicted as presence or background by our environmental variables. While there are many approaches used in species distribution modeling, generalized linear modeling is appropriate for niche description and hypothesis testing and is preferred when studying habitat associations at larger spatial scales or when there is considerable sampling bias [33].

We tested the predictive ability of the models by computing the Area Under the Curve (AUC) of the Receiver Operating Characteristic curve values for each model using the *dismo* package in R [31]. An AUC value of 0.5 indicates that the model has no discriminatory ability, while an AUC value of 1 indicates that the model can perfectly discriminate between Golden-winged Warbler presence and absence. We predicted suitable habitat across the landscape of the study area using the *predict* function from the *raster* package, which calculates a suitability index value for every pixel on the landscape. We back-transformed the prediction raster using the inverse logit function, resulting in suitability index values on a scale from 0 to 1.

We created suitability maps for eBird and Audubon North Carolina datasets in R using the transformed prediction raster [31]. For a more accurate visual representation of the data, we created breaks in the suitability index color schemes at 10 class intervals for each dataset using the quantile method.

## Results

Model results from the Audubon NC dataset indicate that at the local scale (within 150 m) Golden-winged Warbler presence was positively associated with percent forest of height 0-10 meters (P = 1.20e-08), negatively associated with percent forest of height 25-50 meters (P = 1.26e-05), and positively associated with percent road cover (P < 2e-16) (Table 1). At the landscape scale (within 2.5 km), presence was negatively associated with percent road cover (P = 2.33e-13) (Table 1). The Audubon model received an AUC value of 0.75.

**Table 1.** Global linear model results for Audubon NC and eBird datasets including variables at local (150m) and landscape (2.5km) scales.

|  | Audubon model |  | eBird model |  |
| --- | --- | --- | --- | --- |
| Variable | Estimate | Std. Error | Estimate | Std. Error |
| (Intercept) | -2.44*** | 0.23 | -3.65*** | 0.38 |
| Forest height 0-10m within 150m | **7.14\*\*\*** | 1.25 | **4.59\*** | 2.03 |
| Forest height 25-50m within 150m | **-2.34\*\*\*** | 0.53 | **-5.69\*\*\*** | 1.25 |
| Herb/shrub within 150m | -0.04 | 0.64 | -0.96 | 1.09 |
| Road cover within 150m | **13.34\*\*\*** | 1.38 | **12.19\*\*\*** | 1.88 |
| Forest height 25-50m within 2.5km | -0.20 | 0.83 | -0.45 | 1.37 |
| Road cover within 2.5 km | **-50.03\*\*\*** | 6.83 | **-28.36\*\*\*** | 8.16 |
| Developed land within 2.5 km | -50.66 | 27.11 | 9.90 | 8.52 |

Model results using eBird data were remarkably similar compared to those using the Audubon NC dataset. At the local scale, presence was positively associated with percent forest of height 0-10 meters (P = 0.023), negatively associated with percent forest of height 25-50 meters (P = 5.62e-06), and positively associated with percent road cover (P = 9.11e-11) (Table 1). At the landscape scale (within 2.5 km), presence was negatively associated with percent road cover (P = 0.00051) (Table 1). The eBird model received an AUC value of 0.80.

Habitat suitability maps created for each dataset show that most of the landscape is not well-suited for breeding Golden-winged Warblers based on the models (Fig 2). Mean suitability values across the landscape of the study area were only 0.026 (standard deviation, 0.032) based on the Audubon model and 0.0085 (standard deviation, 0.014) based on the eBird model. The eBird and Audubon prediction rasters were positively correlated (Spearman’s correlation coefficient = 0.65, P < 2e-16).

**Fig 2.**
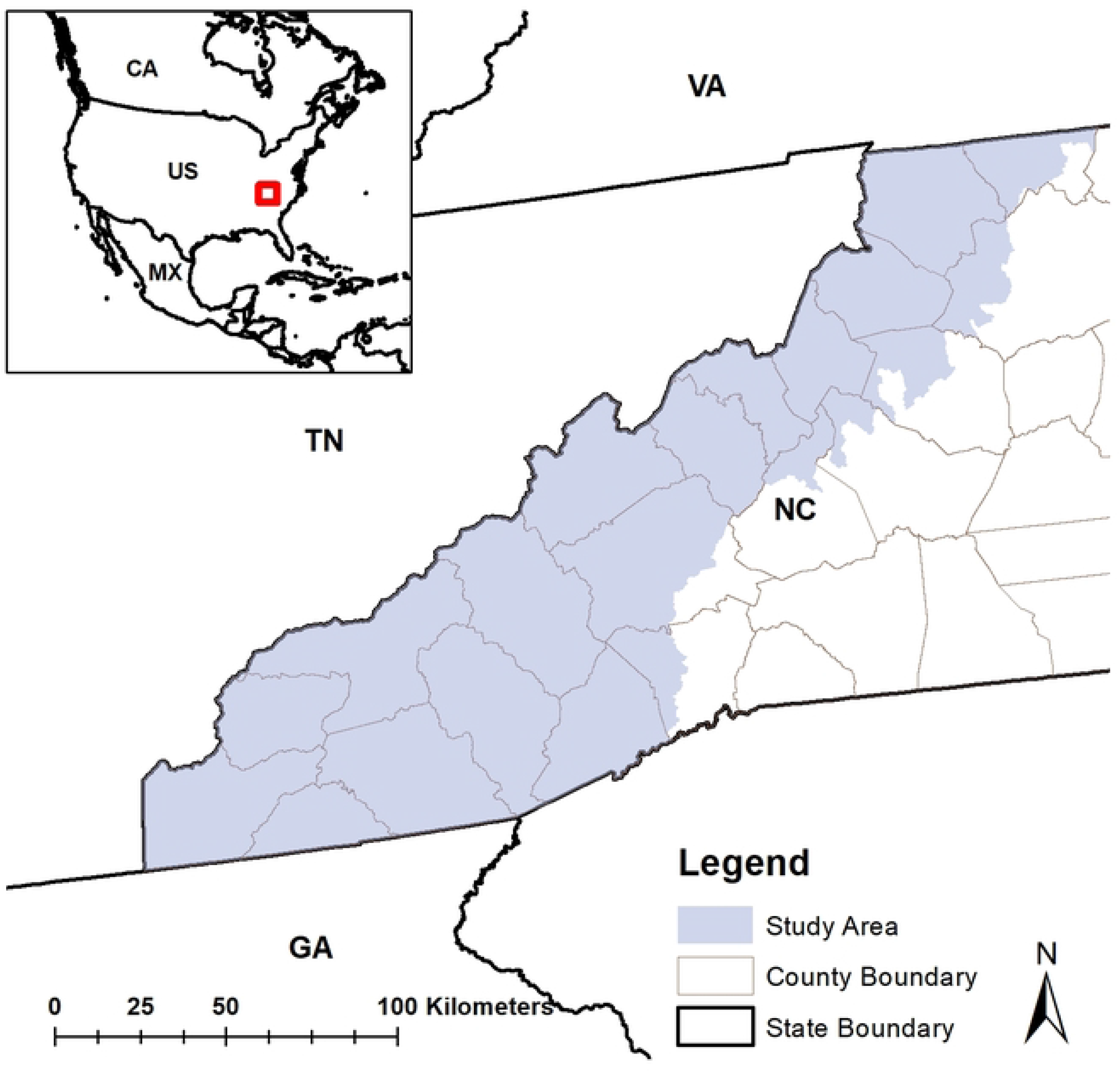
Habitat suitability prediction maps created from (A) Audubon and (B) eBird models.

## Discussion

We predicted habitat suitability across Western North Carolina using community science data from eBird and structured survey data from Audubon NC. Our results suggest that in Western North Carolina, Golden-winged Warblers are found at sites with less mature forest, more young forest, and more road cover at a local scale (within 150m). At a landscape scale (within 2.5 km), Golden-winged Warblers prefer less road cover (Table 1). These findings demonstrate the importance of considering land use variables at different spatial scales when studying Golden-winged Warbler habitat, since different variables are important at different scales and variables may have opposite effects depending on scale. Notably, Audubon and eBird models identified the same significant predictor variables with similar parameter estimates on the same order of magnitude. In the following discussion of our results, we outline the similarities and differences between our models and the Golden-winged Warbler Working Group (GWWG) management guidelines, provide management recommendations based on our results, and conclude with further applications of our work.

Our results are mostly compatible with GWWG management guidelines with a few caveats. GWWG management guidelines call for >70% forest cover within 2.4km of a habitat patch and 60-80% forest cover within 240m of a habitat patch [20]. Our results suggest that in Western North Carolina, mature forest cover at a landscape scale is not an important predictor of suitable Golden-winged Warbler habitat (Table 1). Most likely, this result is due to the landscape being dominated by mature forest cover, so percent mature forest cover is not important in distinguishing background points from presence points. Additionally, the GWWG recommends 15-55% shrub-herbaceous cover within 240m of a habitat patch [20]. Percent herb/shrub at the local scale was not a significant predictor of presence in our models. However, this could be reflective of the nature of the LandFire dataset to underrepresent early successional habitat. Notably, GWWG management guidelines suggest maintaining 30-70% shrub and sapling cover within a habitat patch [20], which aligns well with our models that show a positive association between presence and percent young forest at a local scale (Table 1). Considering the significance of young forest as a predictor, it is likely that in Western North Carolina, young forest accounts for most of the early successional habitat in the LandFire dataset, rather than herb/shrub cover. Finally, GWWG management guidelines indicate that developed land is unsuitable for breeding Golden-winged Warblers [20]. Although our models show no effect of the developed land LandFire category, we did find a significant effect of road cover at both spatial scales (Table 1). Road cover at the landscape scale is negatively associated with presence, indicating that Golden-winged Warblers are selecting habitat within a landscape that is less developed. Road cover at the local scale is positively associated with presence, but this can be attributed to 1) the propensity for early-successional habitat to be near roads; and 2) surveyor bias due to accessibility.

It is important to note that the GWWG outlines management goals to create ideal Golden-winged Warbler breeding habitat. In reality, Golden-winged Warblers may be selecting sites that are less than optimal based on what is available. For example, the lack of early successional habitat in our study area may force Golden-winged Warblers to use very small corridors of early successional habitat with unmeasurable (with spatial data) amounts of herb/shrub cover. Thus, differences between our model results and the GWWG recommendations do not disqualify those recommendations, but rather describe how habitat is being used in Western North Carolina in contrast to those recommendations.

Based on our results, we make the following management and conservation recommendations for the Western North Carolina subregion. Both the Audubon and eBird models identified road cover within 2.5km as the most important predictor of Golden-winged Warbler presence based on the magnitude of the parameter estimates. This suggests that in Western North Carolina, road density at the landscape scale is driving the difference between background sites and Golden-winged Warbler breeding sites. We recommend that Golden-winged Warbler management efforts be concentrated on areas of the landscape with low road density at the landscape scale (2.5km). Both models also found young forest (0-10m in height) to be a better predictor of Golden-winged Warbler presence than herb/shrub cover at the local scale. The young forest LandFire category likely signals for vegetation layer complexity, since high light levels in an open canopy of young trees allow growth of herb/shrub vegetation. In our study area, herb/shrub communities without young trees are likely maintained through heavy human disturbance. Thus, Golden-winged Warblers are likely selecting early successional habitat with complex vegetation layers including herbaceous, shrub, and young trees, and with relatively low human disturbance, which is consistent with the literature [10–15]. We recommend that local-scale habitat (150m) be maintained such that young trees are present but space between trees and open canopy allow herb/shrub communities to coexist with young forest.

Both models had AUC values of ≥ 0.75, indicating that the models performed well but there were discrepancies in the data that could not be explained by our predictor variables. Much of this variance can likely be explained by (1) the coarseness and quality of LandFire data and (2) the rareness of Golden-winged Warblers across our study area. The LandFire dataset is an important tool used frequently in spatial ecology, but it has notable shortcomings, including coarse pixel size, limited ground-truthing, and limited ability to describe heterogeneous landscapes. Additionally, LandFire data provides a snapshot of a landscape in time and seasonal variation and ecological succession complicate LandFire’s ability to describe a habitat. The scarcity of Golden-winged Warblers presents a challenge when studying breeding habitat associations since their low abundance can be due to a variety of factors not related to availability of breeding habitat, including dispersal effects, availability and quality of wintering habitat, and migration routes and availability of stopover sites. All these factors can affect the abundance and spatial distribution of Golden-winged Warblers across the landscape of our study area. While our model predictions suggest low habitat suitability across our study area, there could be other factors contributing to the density and distribution of the species across the study area, and these unknown factors could help explain discrepancies in the data that are not explained by the models.

Future research on the habitat associations of Golden-winged Warblers should focus on analysis of multiple sources of spatial data, perhaps of finer resolution, at multiple spatial scales. Since early successional habitat transforms over time into mature forest in our study area, models including land use change with time could help account for habitat succession over time, which we did not account for in this study. Our results show that eBird data can produce species distribution modeling results that are similar to results obtained from more standardized survey methods. Researchers should continue to utilize eBird data to answer ecological questions since eBird data tends to be more comprehensive across both space and time than other methods of data collection. Additionally, since structured surveys such as Audubon North Carolina surveys require time and financial resources and may create a higher level of disturbance from playback, eBird data should be considered as a viable alternative to traditional surveys when appropriate. However, increased detection probability with playback and the increased sample size of occurrences as a result underscore the importance of continuing to use structured survey protocols such as those used in the Golden-winged Warbler Atlas Project when necessary. At the least, eBird and more traditional survey methods should be used in combination or to supplement each other to improve our understanding of Golden-winged Warbler distribution.

## Acknowledgements

We thank Audubon North Carolina and Curtis Smalling for collecting and sharing Golden-winged Warbler Atlas Project data with us. We thank Highlands Biological Station and James T. Costa for connecting Audubon North Carolina (and Aimee Tomcho) with Lauren Whitenack through the UNC-Chapel Hill Institute for the Environment Highlands Field Site student internship program. Lauren Whitenack and Allen Hurlbert conceived the idea for this paper and designed the experiment. Lauren Whitenack analyzed the data in R and wrote the manuscript. Sara Snell Taylor provided mentorship throughout the project, helped design the methods, and helped create all figures. Aimee Tomcho contributed valuable subject matter expertise. All coauthors helped edit and revise the manuscript. Data were collected by Audubon North Carolina employees and volunteers as well as eBird contributors.

## Notes

### Competing Interest Statement

The authors have declared no competing interest.

